# Mechanisms of peptide agonist dissociation and deactivation of adhesion G-protein-coupled receptors

**DOI:** 10.1101/2024.09.07.611823

**Authors:** Keya Joshi, Yinglong Miao

## Abstract

Adhesion G protein–coupled receptors (ADGRs) belong to Class B2 of GPCRs and are involved in a wide array of important physiological processes. ADGRs contain a GPCR autoproteolysis-inducing (GAIN) domain that is proximal to the receptor N-terminus and undergoes autoproteolysis during biosynthesis to generate two fragments: the N-terminal fragment (NTF) and C-terminal fragment (CTF). Dissociation of NTF reveals a tethered agonist to activate CTF of ADGRs for G protein signaling. Synthetic peptides that mimic the tethered agonist can also activate the ADGRs. However, mechanisms of peptide agonist dissociation and deactivation of ADGRs remain poorly understood. In this study, we have performed all-atom enhanced sampling simulations using a novel Protein-Protein Interaction-Gaussian accelerated Molecular Dynamics (PPI-GaMD) method on the ADGRG2-IP15 and ADGRG1-P7 complexes. The PPI-GaMD simulations captured dissociation of the IP15 and P7 peptide agonists from their target receptors. We were able to identify important low-energy conformations of ADGRG2 and ADGRG1 in the active, intermediate, and inactive states, as well as exploring different states of the peptide agonists IP15 and P7 during dissociation. Therefore, our PPI-GaMD simulations have revealed dynamic mechanisms of peptide agonist dissociation and deactivation of ADGRG1 and ADGRG2, which will facilitate rational design of peptide regulators of the two receptors and other ADGRs.

## INTRODUCTION

G protein-coupled receptors (GPCRs) are critical membrane proteins involved in various physiological functions, including vision, neurotransmission, endocrine, and immune response(*1, 2*). Due to the critical roles in cellular signaling, approximately 34% of FDA-approved therapeutic agents act on GPCRs, providing an important framework for drug design of GPCRs(*3–5*). Adhesion GPCRs (ADGRs) play key roles in tissue development and regulation of the reproductive, nervous, cardiovascular and endocrine systems(*6–11*). ADGRG1 or GPR56 is widely distributed and implicated in immune system functions, brain development, and male fertility(*12, 13*). Dysregulation of ADGRG1 is associated with cancer(*13–17*) and cortical brain malformation disorders, including bilateral frontoparietal polymicrogyria(*18*). ADGRG2 plays an important regulatory role in male fertility(*19, 20*). Patients suffering from medical condition called congenital bilateral absence of vas deferens reported a hemizygous loss-of-function mutation (c.G118T: p.Glu40*), which leads to an early translational termination in the third exon of ADGRG2(*20, 21*).

Previous studies have reported the activation of ADGRG2 in the presence of tethered peptide agonist p15 with sequence “TSFGILLDLSRTSLP”(*22*). However, there is low binding affinity reported of p15 for ADGRG2. Sun et al. addressed the low binding affinity of p15 by designing an optimized peptide agonist VPM-15 by mutating the residues in the Stachel sequence. The first residue, which is threonine (T) was mutated to valine (V) and the third residue, phenylalanine (F) was mutated to 4-methyl phenylalanine (4PH) to produce an optimized peptide agonist VPM-15 with sequence “VS4PHGILLDLSRTSLP”(*22*). With the introduction of these mutations, the binding affinity improved significantly compared to the tethered peptide agonist p15(*22*). In a later study, Xiao et al. generated a new optimized peptide agonist IP15 by mutating the first residue threonine of VPM-15 to isoleucine. Due to this mutation, there was a 10,000-fold increase in the peptide binding affinity. This modification also led to the cryo-EM structure of ADGRG2-IP15 in complex with the Gs protein (PDB:7WUI)(*23*). Recently, our lab performed all-atom Gaussian accelerated Molecular Dynamics (GaMD) simulations using the cryo-EM structure of the ADGRG2–IP15-Gs complex (PDB: 7WUI)(*23*), which revealed distinct binding conformations of the agonist and antagonist peptides in ADGRG2(*24*).

In 2022, a cryo-EM structure was reported for the ADGRG1 in complex with the G_13_ protein (PDB:7SF8)(*25*). The first 7 amino acids with sequence “TYFAVLM” act as a tethered peptide agonist (P7) of ADGRG1. In this configuration, the P7 stalk peptide bends nearly 180° downward into the orthosteric site and forms important interactions. One such interaction is formed with the extracellular loop 2 (ECL2), which reaches into the interior of the orthosteric pocket to form a wedge-like plug structure. Residue W557^45.51^ in ECL2 interacts with residue L388 in the tethered peptide agonist (p7)(*25*). Notably, W421^45.51^ in the ECL2 of GPR97 also reaches down into the orthosteric pocket close to the bound glucocorticoid ligand(*26*).

Building on GaMD(*27–32*), a new PPI-GaMD approach has been developed to explore the PPIs, in which the interaction energy potential of protein binding partners (both electrostatic and van der Waals interactions) is selectively boosted to enhance protein dissociation. In addition, another boost is simultaneously applied on the remaining potential energy of the system to enhance the rebinding of the proteins(*33*). Here, we have applied PPI-GaMD simulations to elucidate mechanisms of the dissociation of the peptide agonists and deactivation of the ADGRs. We have used PPI-GaMD method for this study instead of the Peptide GaMD (Pep-GaMD) method (*34*) as the peptide agonists IP15 and P7 have very high binding affinities for the protein and can be considered as small protein molecules for simulations. With PPI-GaMD(*33*), dissociation of the peptide agonists was efficiently accelerated by boosting the interaction energy potential between the two partners, i.e., the receptor and peptide agonist.

## METHODS

### System Setup

The cryo-EM structures of the active IP15-Gs-bound ADGRG2 (PDB:7WUI) and active P7-G_13_-bound ADGRG1 (PDB:7SF8) complexes were used to set up the ADGRG2 and ADGRG1 simulation systems, respectively (*23, 25*). Missing residues in the extracellular loop 2 (ECL2) and intracellular loop 3 (ICL3) were added to the ADGRG2-IP15 complex using the Swiss Modeler(*35*). The last four missing residues in the IP15 agonist (TSLP) were added by copying their coordinates from the 7WUQ PDB structure (*23*). The first 7 amino acids with sequence “TYFAVLM” act as a tethered peptide agonist (P7) for ADGRG1(*25*). The covalent bond between the residues M389 in the tethered peptide agonist (P7) and V390 in the linker was manually removed to obtain the peptide agonist P7. Missing residues in the extracellular loop 2 (ECL2) and helix 8 (H8) were also added to the ADGRG1-P7 complex using the Swiss Modeler (*35*). G protein was removed to obtain only the agonist IP15-bound ADGRG2 and P7-bound ADGRG1 structures **(Figure S1A and S4A)**.

The peptide-bound ADGRG2 and ADGRG1 receptors were prepared and embedded in a phosphatidylcholine (POPC) lipid bilayer using the CHARMM-GUI online server (*36–41*). The residues at the protein termini were assigned neutral patches (acetyl and methyl amide). The peptide termini were kept charged (NH3+ and COO−). The system was neutralized by adding counter ions and immersing in a cubic TIP3P waterbox (*42*) which was extended for 25 Å from the peptide−protein complex surface **(Figure S1B and S4B)**. The CHARMM36m (*43*)force-field parameters were used, and the CHARMM-GUI output files and scripts were used for PPI-GaMD simulations (*33*).

ADGRG2–IP15 system measured about 80.26 × 80.26 × 133.17 Å^3^ with 152 POPC lipid molecules (77 molecules on the upper leaflet and 75 molecules on the lower leaflet) and 18,188 water molecules while ADGRG1-P7 system measured 90.00 × 90.00 × 124.66 Å^3^ with 152 POPC lipid molecules (76 molecules on the upper leaflet and 76 molecules on the lower leaflet) and 16,422 water molecules.

### Simulation Protocol

A total of 5000 steps of energy minimizations were carried out on the ADGRG2 and ADGRG1 systems, and a constant number, volume, and temperature (NVT) ensemble equilibration was performed for 125 ps at 310 K. Using an NPT ensemble, additional equilibration was carried out for 375 ps at 310 K. We then performed conventional MD (cMD) simulations on the systems for 40 ns at 1 atm pressure and 310 K temperature with the AMBER22 software package (*44*). Long-range electrostatic interactions were computed with the particle mesh Ewald summation method, and a cutoff distance of 9 Å was used for the short-range electrostatic and van der Waals interactions (*45*). After the cMD runs, we performed PPI-GaMD equilibration for 60 ns for the ADGRG2–IP15 and ADGRG1-P7 complex systems.

In order to observe peptide agonist dissociation during the PPI-GaMD production while keeping the boost potential as low as possible for accurate energetic reweighting, the (σ0P, σ0D) parameters were set to (1.7, 6.0 kcal/mol) and (1.9, 6.0 kcal/mol) for the ADGRG2 and ADGRG1 systems, respectively. This was followed by eight independent PPI-GaMD production runs for 1500 ns for the agonist-bound ADGRG2 and ADGRG1 systems. PPI-GaMD production simulation frames were saved every 0.2 ps for analysis. The PPI-GaMD simulations on the ADGRG2 and ADGRG1 systems are summarized in **Supplementary Tables 1 and 2.**

### Simulation Analysis

Simulation analysis was carried out using CPPTRAJ (*46*) and VMD (*47*). The software tools were applied to track the dissociation of peptide agonists from the ADGRG2 and ADGRG1 receptors. Root mean square deviation (RMSD) of the IP15 agonist and L723^3.58^-R833^6.40^ distance were selected to calculate the 2D free energy profiles to characterize the peptide agonist dissociation and deactivation of ADGRG2. Similarly, RMSD of the P7 agonist and V504^3.58^-L610^6.40^ distance were selected to calculate the 2D free energy profiles to characterize the peptide agonist dissociation and deactivation of ADGRG1. 2D free energy profiles were reweighted using the PyReweighting toolkit (*48*). A bin size of 2 Å was used for the TM3-TM6 distance and 5 Å for RMSDs of the IP15 and P7 agonists. The cutoff was set to 500 frames in one bin for reweighting. The Dpeak clustering algorithm (*49*) was used to cluster snapshots of the receptor and peptide conformations with all PPI-GaMD production simulations combined for each system. The PPI-GaMD simulation snapshots for the IP15 and P7 dissociation from the ADGRG2 and ADGRG1 were clustered, respectively, to identify the representative low-energy conformations in the 2D free energy profiles (**Figure 1D and 3D**).

**Figure 1.**
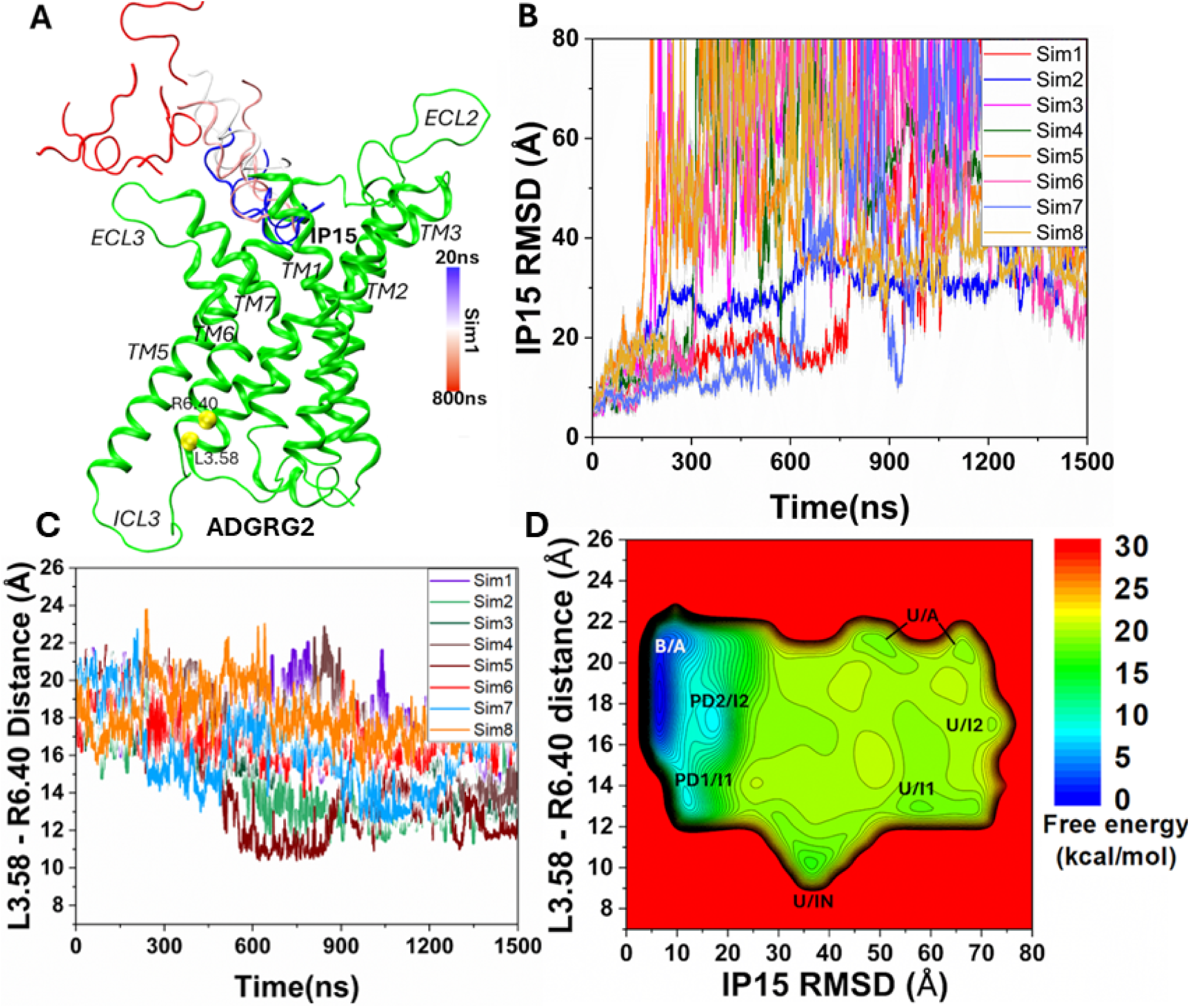
Dissociation of the IP15 peptide agonist and deactivation of the human ADGRG2 receptor observed in the PPI-GaMD simulations: **(A)** A representative dissociation pathway of the IP15 peptide agonist. **(B)** Root-mean-square deviations (RMSDs) of the IP15 peptide agonist relative to the cryo-EM bound conformation (PDB: 7WUI) calculated from the eight 1500 ns PPI-GaMD simulations. **(C)** The TM3-TM6 distance between the Cα atoms of L^3.58^ and R^6.40^ calculated from the eight 1500ns PPI-GaMD simulations. **(D)** 2D potential of mean force (PMF) free energy profile of the IP15 RMSD and L^3.58^-R^6.40^ distance calculated by combining the eight PPI-GaMD simulations. The low-energy states are labeled as “Bound/Active” (B/A), “Partially Dissociated 1/Intermediate 1” (PD1/I1), “Partially Dissociated 2/Intermediate 2” (PD2/I2), “Unbound/Active” (U/A), “Unbound/Intermediate 1” (U/I1), “Unbound/Intermediate 2” (U/I2) and “Unbound/Inactive” (U/IN).

## RESULTS AND DISCUSSION

### PPI-GaMD Simulations Captured Peptide Agonist Dissociation from the ADGRG2 Receptor

In eight independent 1500ns PPI-GaMD production simulations, the IP15 peptide agonist dissociated from the orthosteric site of the ADGRG2 receptor at ∼200-800ns, for which RMSD of the peptide relative to the cryo-EM bound conformation increased to ≥ 30 Å (**Figure 1A and 1B**). In all PPI-GaMD simulations, IP15 dissociated completely into the solvent from the ADGRG2 following a similar pathway through the extracellular mouth between ECL2, ECL3, TM7 and TM1 (**Figure 1A**). The eight PPI-GaMD simulations showed similar boost potentials with averages of ∼28.6-31.7 kcal/mol and standard deviations (SDs) of ∼4.4-5.4 kcal/mol **(Table S1)**.

We combined all the PPI-GaMD production simulations to calculate free energy profiles for dissociation of the IP15 agonist from the ADGRG2 through energetic reweighting **(See Methods).** In the 2D free energy profile of the IP15 agonist RMSD and L723^3.58^-R833^6.40^ distance (**Figure 1D**), we identified seven low-energy conformational states, i.e., the “Bound/Active” (B/A), “Partially Dissociated 1/Intermediate 1” (PD1/I1), “Partially Dissociated 2/Intermediate 2” (PD2/I2), “Unbound/Active” (U/A), “Unbound/Intermediate 1” (U/I1), “Unbound/Intermediate 2” (U/I2) and “Unbound/Inactive” (U/IN).

### Low-energy conformational states of IP15 Dissociation from the ADGRG2 Receptor

The Density Peak (DPeak) clustering algorithm (*49*) was used to cluster snapshots of the ADGRG2 with all the PPI-GaMD production simulations combined to identify the representative low-energy conformations of the IP15-ADGRG2 system (**Figures 1 and 2**). In the “Bound/Active” (B/A) state, RMSD of the IP15 agonist centered around ∼5 Å and the distance between receptor residues L723^3.58^ and R833^6.40^ centered around ∼20 Å, being similar to the cryo-EM structure (**Figure 1D**). The peptide adopted an alpha-helical conformation, stabilized by receptor residues from TM1, TM2, TM5, TM6, TM7, ECL3 and ECL2 **(Figure S2)**. The alpha-helical conformation of IP15 agonist showed a clear division of hydrophilic residues “- - - - - - - SRTS - -” on the upper rim and hydrophobic residues “TSFGILLDL - - - - LP” on the lower rim. The lower rim of the peptide agonist assumed an alpha-helical conformation, deeply inserted into the hydrophobic core of the receptor and showed interactions with receptor residues Y633^1.44^, F679^2.64^, W779^ECL2^, Y787^5.36^, F850^6.57^ and F864^7.42^ **(Figure S2B)**. Previous studies have experimentally validated W779^ECL2^ residue in ECL2 to be critical for the activation of ADGRG2 by the Stachel sequence (*50*). The IP15 peptide agonist could not activate the ADGRG2 receptor when W779^ECL2^ was mutated to alanine. The upper rim of the IP15 agonist comprised of hydrophilic residues in which R609 formed a polar interaction with receptor residue Y787^5.36^ **(Figure S2B).** Ala substitution of Y787^5.36^ in ADGRG2 significantly impaired the constitutive activity of ADGRG2 (*50*).

**Figure 2.**
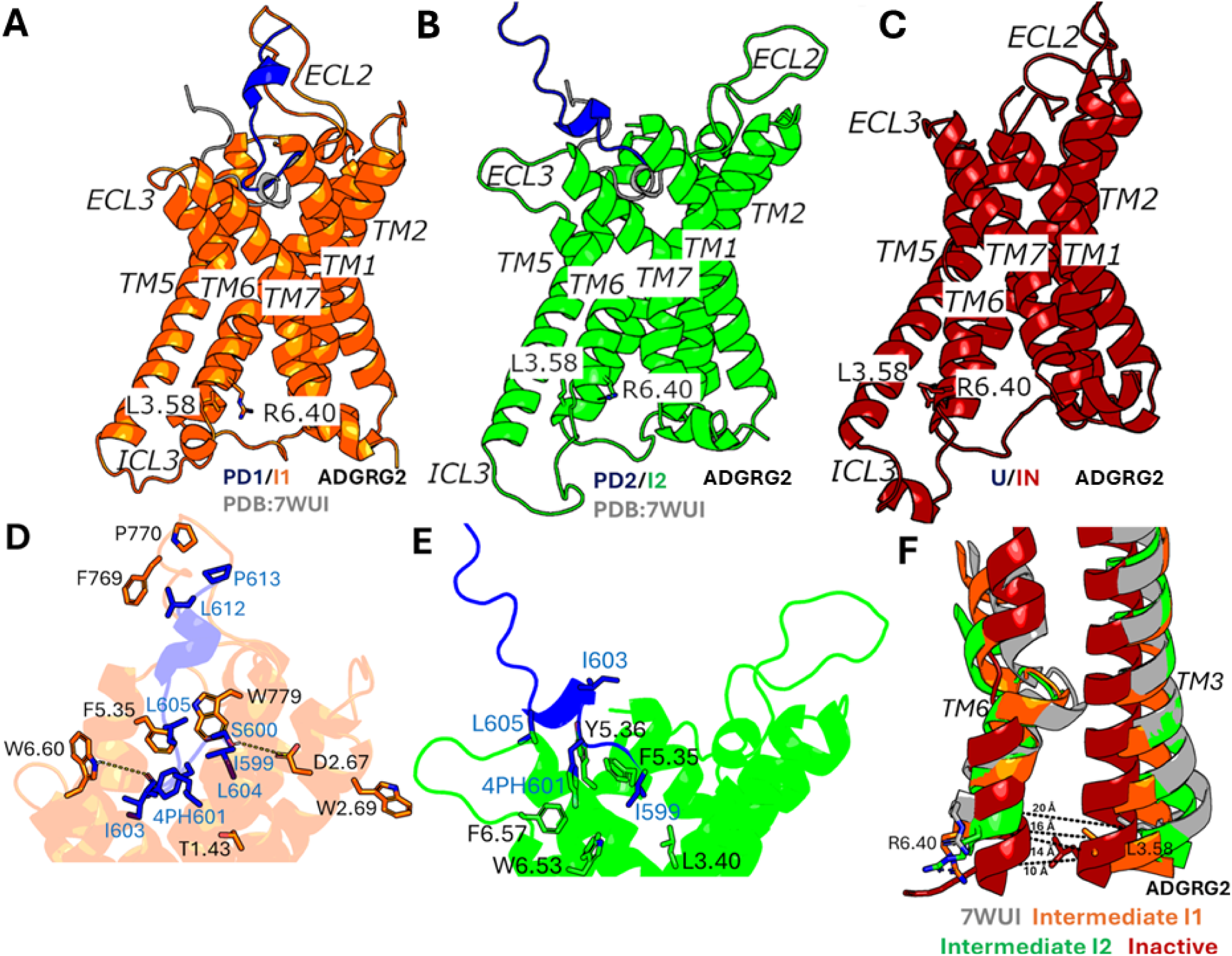
Low-energy conformational states of the IP15 peptide dissociation from the ARGRG2 receptor: **(A-B)** The “Partially Dissociated 1/Intermediate 1” (PD1/I1) and “Partially Dissociated 2/Intermediate 2” (PD2/I2) states. The cryo-EM bound conformation (PDB: 7WUI) is shown in grey as reference. **(C)** The “Unbound/Inactive” (U/IN) state. **(D)** Critical interactions between IP15 (blue) and ADGRG2 (orange) observed in the (PD1/I1) state. The peptide agonist formed hydrogen-bonding (yellow dash lines) and hydrophobic interactions with receptor residues T^1.43^, D^2.67^, W^2.69^, F^5.35^, F769^ECL2^, P770^ECL2^, W779^ECL2^ and W^6.60^. **(E)** Critical interactions between IP15 (blue) and ADGRG2 (green) observed in the (PD2/I2) state. The peptide agonist formed mostly hydrophobic interactions with receptor residues L^3.40^, F^5.35^, Y^5.46^, W^6.53^ and F^6.57^. **(F)** Unwinding of kink in TM6 observed in Intermediate “I1” (orange), Intermediate “I2” (green) and Inactive (red) states compared with the active cryo-EM conformation (grey).

In the “Partially Dissociated 1/Intermediate 1” (PD1/I1) state, the C-terminus of the IP15 peptide agonist residues (L612 and P613) formed close interactions with receptor residues F769^ECL2^, P770^ECL2^ and W779^ECL2^ (**Figure 2A and 2D**). The N-terminus of the IP15 peptide agonist formed hydrophobic interactions with receptor residues T632^1.43^, W684^2.69^ and F786^5.35^ in the orthosteric pocket. Residues S600 and I603 in the IP15 agonist also formed polar interactions with receptor residues D682^2.67^ and W853^6.60^ (**Figure 2A and 2D**). In the PD1/I1 state, the distance between receptor residues L723^3.58^-R833^6.40^ decreased to ∼14 Å (**Figure 1D and 2F**).

In the “Partially Dissociated 2/Intermediate 2” (PD2/I2) state, the IP15 peptide agonist was located between the receptor ECL3, TM5, TM6 and TM7. In the PD2/I2 conformation, the C-terminus of the IP15 peptide agonist showed zero contacts with the receptor residues while the N-terminus of the IP15 peptide formed hydrophobic interactions with receptor residues L705^3.40^, F786^5.35^, Y787^5.36^, F850^6.57^ and W846^6.53^ in the orthosteric pocket (**Figure 2B and 2E**). Residues 4PH601 and L605 in the IP15 peptide agonist were in direct contact with the putative toggle switch W846^6.53^, which is known to be the activation hub of many GPCRs. Alteration of the conformation of W846^6.53^ could directly modulate ADGRG2 activity (*50*). In the PD2/I2 state, the distance between receptor residues L723^3.58^-R833^6.40^ changed to ∼16 Å (**Figure 1D and 2F**). In the “Partially Dissociated 2” (PD2) state, the ECL2 attained a complete open conformation and showed no interaction with the IP15 peptide agonist while the ECL3 facilitated the peptide dissociation. The IP15 agonist RMSD and distance between residues W779^ECL2^-W853^6.60^ (ECL3) were selected to calculate the 2D free energy profile to characterize the opening of the ECL3 gate **(Figure S3A).** Upon the IP15 peptide agonist dissociation from the orthosteric site, the ECL3 gate opened completely characterized by outward flipped movement of the sidechain of residue W853^6.60^ in ECL3 and the distance between residues W779^ECL2^ and W853^6.60^ increased to ∼30 Å **(Figure S3B)**.

In the “Unbound/Active” (U/A) state, the IP15 peptide agonist dissociated from the orthosteric site, for which the RMSD of the peptide relative to the cryo-EM bound conformation increased to ≥ 50 Å. In the U/A state, the distance between receptor residues L723^3.58^-R833^6.40^ remained ∼20 Å, conforming to the active state of ADGRG2 (**Figure 1D**).

In the “Unbound/Intermediate 1” (U/I1) and “Unbound/Intermediate 2” (U/I2) states, the IP15 peptide agonist dissociated from the orthosteric site, for which the RMSD of the peptide relative to the cryo-EM bound conformation increased to ≥ 60 Å and ≥ 80 Å, respectively. In the U/I1 state, the distance between receptor residues L723^3.58^-R833^6.40^ decreased to ∼14 Å (**Figure 1D and 2F**), while in the U/I2 state, the distance between receptor residues L723^3.58^-R833^6.40^ decreased to ∼16 Å (**Figure 1D and 2F**).

In the “Unbound/Inactive” (U/IN) state, the IP15 peptide agonist dissociated from the orthosteric site, for which the RMSD of the peptide relative to the cryo-EM bound conformation increased to ≥ 40 Å and the distance between receptor residues L723^3.58^-R833^6.40^ decreased to ∼10 Å. Therefore, the PPI-GaMD enhanced sampling simulations were able to sample the conformational transition of ADGRG2 from the active to inactive state upon the IP15 peptide agonist dissociation (**Figure 2C**). Interestingly, as ADGRG2 attained the inactive state, the characteristic kink in TM6 disappeared along with closing of the extracellular pocket characterized by the inward movement of the residues involved in the extracellular region of TM5, TM6 and TM7 (**Figure 2C and 2F**).

### PPI-GaMD Simulations Captured Peptide Agonist Dissociation from the ADGRG1 Receptor

In eight independent 1500ns PPI-GaMD production simulations, the P7 peptide agonist dissociated similarly from the orthosteric site of the ADGRG1 receptor at ∼200-800ns, for which the RMSD of the peptide relative to the cryo-EM bound conformation increased to ≥ 30 Å (**Figure 3A and 3B**). In all PPI-GaMD simulations, P7 dissociated completely into the solvent from the ADGRG1 following a similar pathway through the extracellular mouth between ECL2, ECL3, TM7 and TM1 (**Figure 3A**). The eight PPI-GaMD simulations showed similar boost potentials with averages of ∼45.4-49.1 kcal/mol and SDs of ∼4.1-6.6 kcal/mol **(Table S2)**.

**Figure 3.**
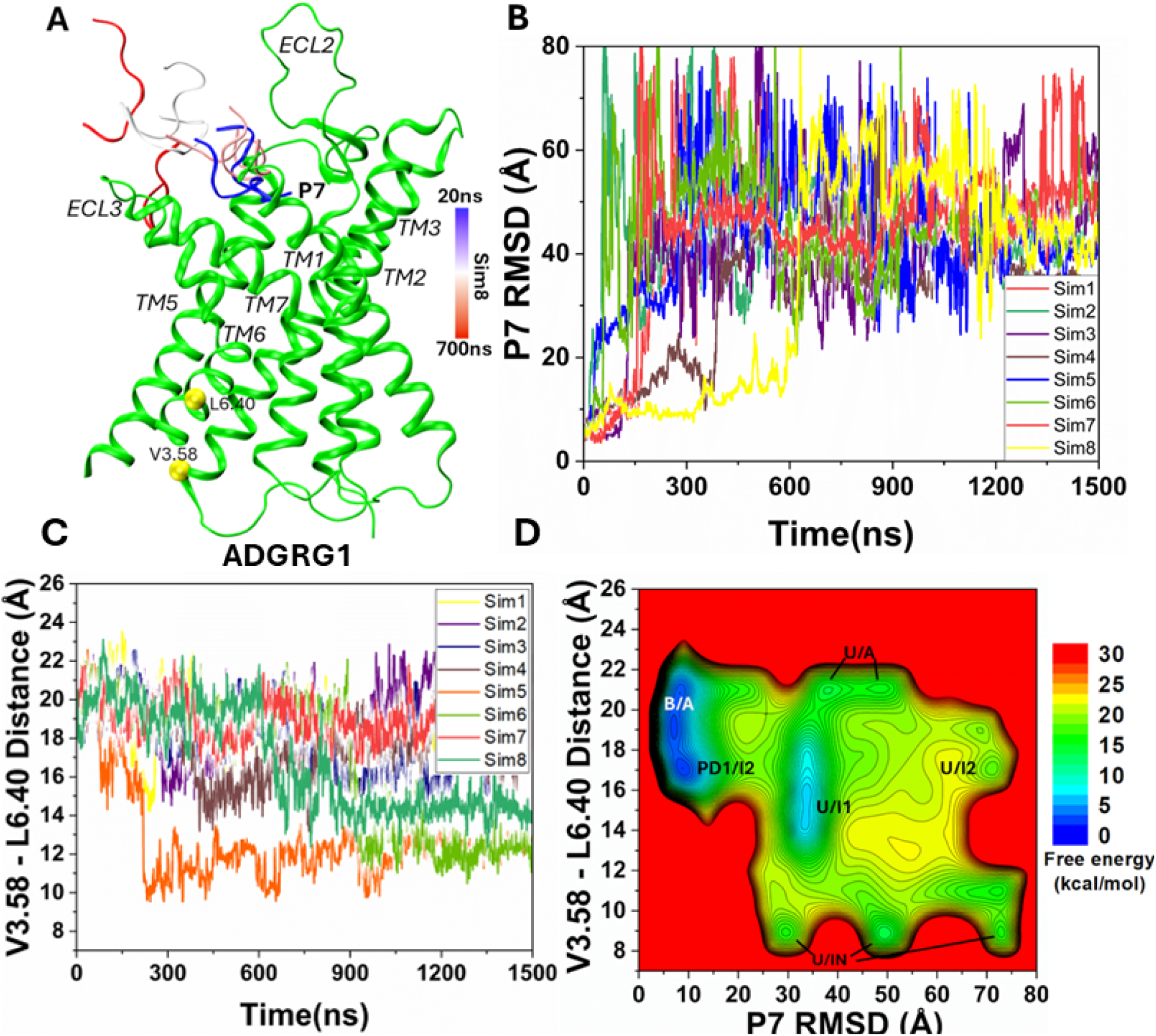
Dissociation of the P7 peptide agonist and deactivation of the human ADGRG1 receptor observed in the PPI-GaMD simulations: **(A)** A representative dissociation pathway of the P7 peptide agonist. **(B)** Root-mean-square deviations (RMSDs) of the P7 peptide agonist relative to the cryo-EM bound conformation (PDB: 7SF8) calculated from the eight 1500 ns PPI-GaMD simulations. **(C)** The TM3-TM6 distance between the Cα atoms of V^3.58^ and L^6.40^ calculated from the eight 1500ns PPI-GaMD simulations. **(D)** 2D potential of mean force (PMF) free energy profile of the P7 RMSD and V^3.58^-L^6.40^ distance calculated by combining the eight PPI-GaMD simulations. The low-energy states are labeled as “Bound/Active” (B/A), “Partially Dissociated 1/Intermediate 2” (PD1/I2), “Unbound/Active” (U/A), “Unbound/Intermediate 1” (U/I1), “Unbound/Intermediate 2” (U/I2) and “Unbound/Inactive” (U/IN).

We also combined all the PPI-GaMD production simulations to calculate free energy profiles for dissociation of the P7 agonist from the ADGRG1. In the 2D free energy profile of the P7 agonist RMSD and V504^3.58^-L610^6.40^ distance (**Figure 3D**), we identified six low-energy conformational states, i.e., the “Bound/Active” (B/A), “Partially Dissociated 1/Intermediate 2” (PD1/I2), “Unbound/Active” (U/A), “Unbound/Intermediate 1” (U/I1), “Unbound/Intermediate 2” (U/I2) and “Unbound/Inactive” (U/IN). Notably, the “Partially Dissociated 2” (PD2) state identified in the IP15-ADGRG2 system (**Figure 1D**), did not appear as a low energy state in the P7-ADGRG1 system simulations.

### Low-energy conformational states of P7 Dissociation from the ADGRG1 Receptor

The DPeak clustering algorithm (*49*) was used to cluster snapshots of the ADGRG1 with all the PPI-GaMD production simulations combined to identify the representative low-energy conformations of the P7-ADGRG1 system (**Figures 3 and 4**). In the “Bound/Active” (B/A) state, RMSD of the P7 agonist centered around ∼5 Å and the distance between receptor residues V504^3.58^ and L610^6.40^ centered around ∼20 Å, being similar to the cryo-EM structure (**Figure 3D**). The peptide bends nearly 180° downward into the orthosteric pocket **(Figure S5)** and showed interactions with receptor residues F454^2.58^, W563^ECL2^, Y571^5.36^, F630^ECL3^, W623^6.53^ and F643^7.42^ **(Figure S5B).** In ADGRG1, the P7 peptide formed interaction with W623^6.53^, a conserved residue that interacts with the bound steroid ligand in the GPR97 partial agonist-activated receptor structure (*26*). The function of the adhesion GPCR W623^6.53^ seems to parallel the ‘toggle switch’ residue W^6.48^ in class A GPCRs which rearranges upon agonist binding and faciliates the opening of the cytoplasmic end of TM6 for G-protein engagement (*51*). Mutation of W623^6.53^ in ADGRG1 (W617A) strongly reduced the receptor-dependent G-protein activation (*25*).

**Figure 4.**
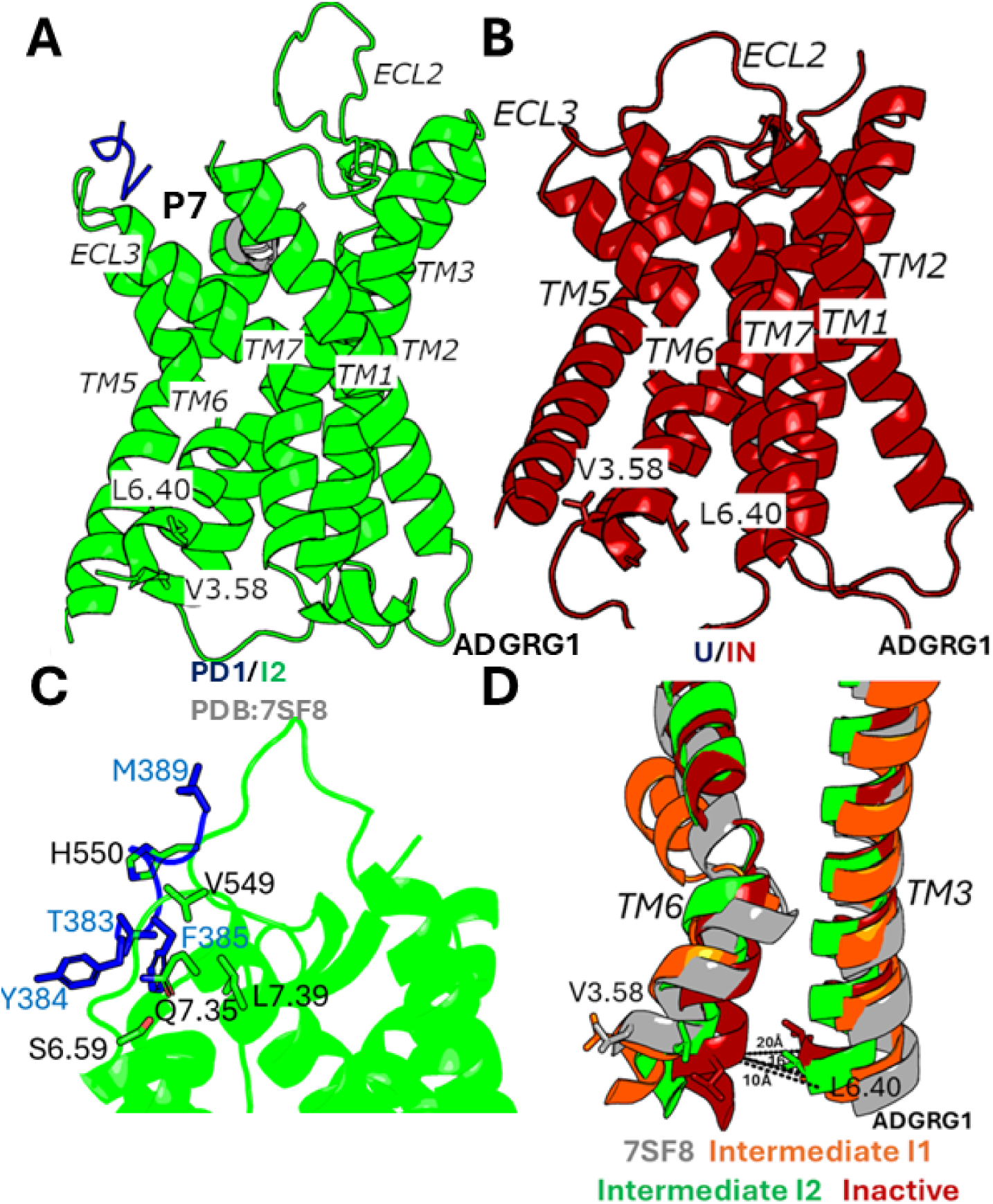
Low-energy conformational state of the P7 peptide dissociation from the ARGRG1 receptor: **(A)** The “Partially Dissociated 1/Intermediate 2” (PD1/I2) state. The cryo-EM bound conformation (PDB: 7SF8) is shown in grey as reference. **(B)** The “Unbound/Inactive” (U/IN) state. **(C)** Critical interactions between P7 (blue) and ADGRG1 (green) observed in the (PD1/I2) state. The peptide agonist formed mostly hydrophobic interactions with receptor residues V549^ECL2^, H550^ECL2^, S^6.59^, Q^7.35^ and L^7.39^. **(D)** Unwinding of kink in TM6 observed in Intermediate “I1” (orange), Intermediate “I2” (green) and Inactive (red) states compared with the active cryo-EM conformation (grey).

In the “Partially Dissociated 1/Intermediate 2” (PD1/I2) state, the P7 peptide agonist formed hydrophobic interactions with receptor residues V549^ECL2^, H550^ECL2^, S629^6.59^, Q636^7.35^ and L640^7.39^ in the orthosteric pocket (**Figure 4C**). In the PD1/I2 state, the distance between receptor residues V504^3.58^-L610^6.40^ decreased to ∼16 Å. (**Figure 4D**). In the “Unbound/Active” (U/A) state, the P7 peptide agonist dissociated from the orthosteric site, for which the RMSD of the peptide relative to the cryo-EM bound conformation increased to ≥ 50 Å. In the U/A state, the distance between receptor residues V504^3.58^-L610^6.40^ remained ∼20 Å (**Figure 4D**).

In the “Unbound/Intermediate 1” (U/I1) and “Unbound/Intermediate 2” states, the P7 peptide agonist dissociated from the orthosteric site, for which the RMSD of the peptide relative to the cryo-EM bound conformation increased to ≥ 40 Å and ≥ 80 Å, respectively. In the U/I1 state, the distance between receptor residues V504^3.58^-L610^6.40^ decreased to ∼14 Å (**Figure 3D**), while in the U/I2 state, the distance between receptor residues V504^3.58^-L610^6.40^ decreased to ∼16 Å (**Figure 4D**).

In the “Unbound/Inactive” (U/IN) state, the P7 peptide agonist dissociated from the orthosteric site, for which the RMSD of the peptide relative to the cryo-EM bound conformation increased to ≥ 30 Å. In the U/IN state, the distance between receptor residues V504^3.58^-L610^6.40^ decreased to ∼10 Å. PPI-GaMD simulations were able to sample the conformational transition of ADGRG1 from the active to fully inactive state upon the P7 peptide agonist dissociation (**Figure 4B**). In (**Figs. 4D and 2F**), the intermediate states seem to show the largest opening of extracellular pocket, for which the inactive state didn’t show significant closing compared with active. However, consistent closing can be described for TM6 in the intracellular pocket.

## CONCLUSIONS

In this study, all-atom PPI-GaMD simulations have been applied to elucidate the mechanisms of peptide agonist dissociation and deactivation of the ADGRs. PPI-GaMD simulations captured distinct low-energy states of peptide agonists IP15 and P7 during dissociation, as well as low-energy conformations of the ADGRG2 and ADGRG1 in the active, intermediate, and inactive states. In class A GPCRs, the distance between the TM3 and TM6 intracellular domains in the inactive conformation is significantly reduced compared to that of the active state (*52, 53*). In the cryo-EM structures of active ADGRG2 (PDB:7WUI) (*23*) and active ADGRG1 (PDB:7SF8) (*25*), the distance between receptor residues L723^3.58^-R833^6.40^ and V504^3.58^-L610^6.40^ was ∼20 Å, which decreased to ∼10 Å upon the peptide agonist dissociation, characterized by the inward movement of TM6 intracellular end and loss of the TM6 kink.

Peptide agonists IP15 and P7 in the “Bound/Active” (B/A) state assumed an alpha-helical conformation, stabilized by residues from the orthosteric pocket **(Figure S2 and S4)**. Mutation of the residues present in the IP15 and P7 peptide agonist or the interacting residues from the orthosteric pocket decreased the receptor’s G protein binding activity (*23, 25*). Mutation of W623^6.53^ and F643^7.42^ to alanine in ADGRG1 decreased the GTP_l1l_S turnover significantly (*25*).

In ADGRG2, the peptide agonist IP15 sampled two different conformations-“Partially Dissociated 1” (PD1) and “Partially Dissociated 2” (PD2) state. In the intermediate states, the peptide agonist IP15 formed important interactions with residues from the orthosteric pocket (**Figure 2D and 2E**). Mutation of the residues F769^ECL2^, Y787^5.36^, F850^6.57^ and W846^6.53^ to alanine showed a direct implication on the cAMP accumulation for the downstream signaling (*23*). In comparison, the peptide agonist P7 in ADGRG1 sampled only the “Partially Dissociated 1” (PD1) state, which was stabilized by hydrophobic interactions with residues V549^ECL2^, H550^ECL2^, S629^6.59^, Q636^7.35^ and L640^7.39^ in the orthosteric pocket (**Figure 4C**). Mutation of the interacting residues or peptide residues to alanine in ADGRG1 decreased the GTP_l1l_S turnover significantly (*25*).

It is also important to mention the opening of the ECL3 gate to facilitate the peptide agonist IP15 dissociation in ADGRG2, characterized by outward flipping of the sidechain of W853^6.60^ residue in ECL3 **(Figure S3B)**. Mutation of W853^6.60^ to alanine decreased the cAMP accumulation in the downstream signaling for ADGRG2 (*23*). We also observed the relaxation and unwinding of TM6 kink in both ADGRG2 and ADGRG1 receptors, which can be a characteristic feature for the deactivation of class B2 adhesion GPCRs (**Figure 2F and 4D**).

In summary, we uncovered a plausible mechanism for the dissociation of the peptide agonists and deactivation in the ADGRs. The understanding we gained regarding deactivation of the two subtypes of ADGRs and the interactions that facilitate the peptide agonist dissociation can provide a valuable foundation for the design and development of novel peptide regulators for ADGRG2/ADGRG1 and other ADGRs.

## ASSOCIATED CONTENT

### Supporting Information

The Supporting Information includes details about the PPI-GaMD method, Tables S1 and S2, and Figures S1 to S5. This information is available free of charge via the Internet at http://pubs.acs.org.

## AUTHOR INFORMATION

### Corresponding Author

**Dr. Yinglong Miao**− Department of Pharmacology and Computational Medicine Program, University of North Carolina – Chapel Hill, Chapel Hill, NC 27599, USA.

### Author

**Keya Joshi**-Department of Pharmacology and Computational Medicine Program, University of North Carolina – Chapel Hill, Chapel Hill, NC 27599, USA.

### Author contributions

Y.M. designed the research; K.J. performed research; K.J. and Y.M. analyzed data; and K.J. and Y.M. wrote the paper.

### Notes

The authors declare no competing financial interest.

## ACKNOWLEDGMENTS

This work used supercomputing resources with allocation award TG-MCB180049 through the Extreme Science and Engineering Discovery Environment ACCESS, which is supported by National Science Foundation grant number ACI-1548562, and project M2874 through the National Energy Research Scientific Computing Center (NERSC), which is a U.S. Department of Energy Office of Science User Facility operated under Contract No. DE-AC02-05CH11231. This work was supported by the National Institutes of Health (R01GM132572) and the startup funding project 27110 at the University of North Carolina-Chapel Hill.

## TOC Graphic

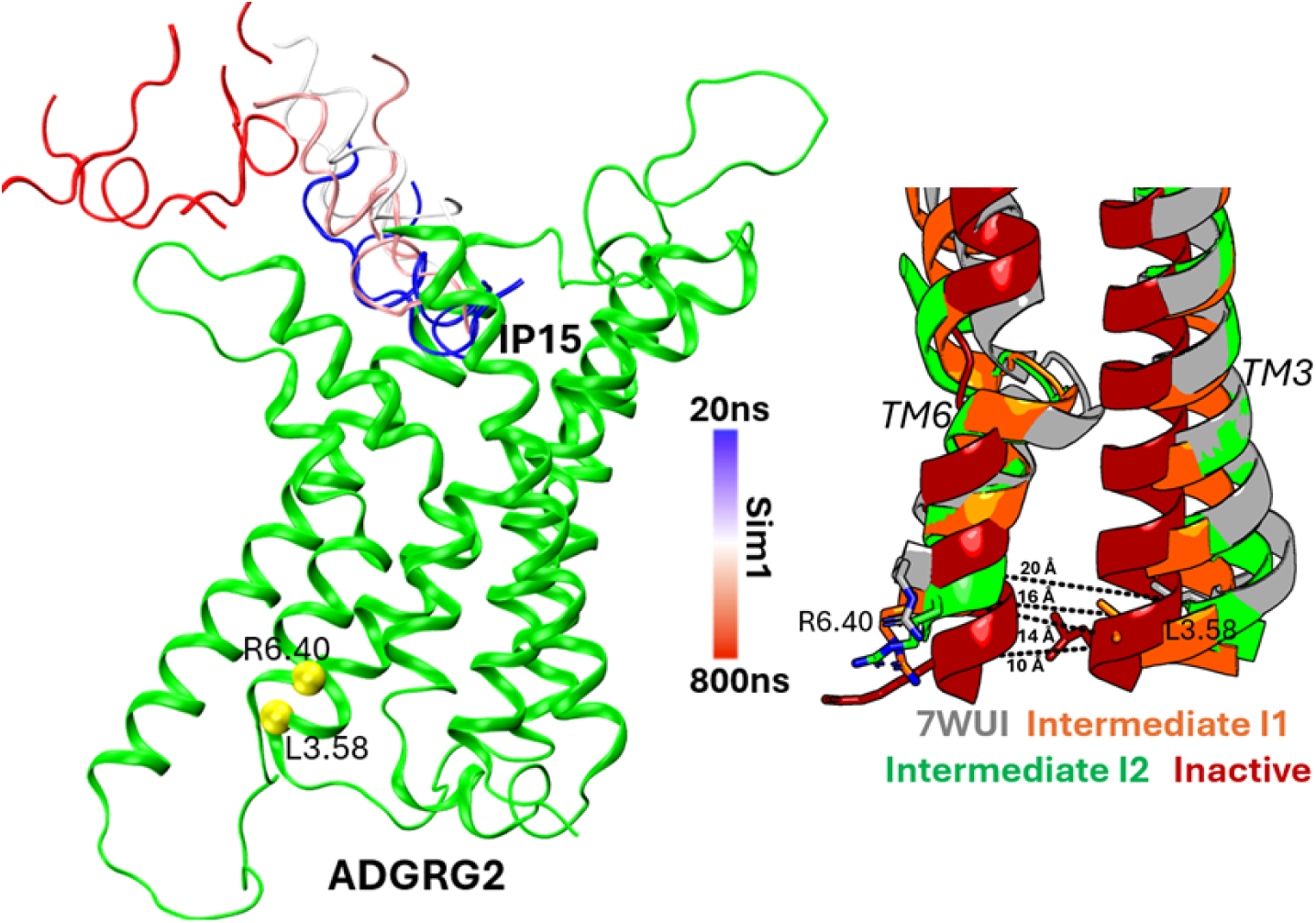

Protein-Protein Interaction-Gaussian accelerated Molecular Dynamics (PPI-GaMD) simulations revealed dynamic mechanisms of peptide agonist dissociation and deactivation of adhesion G protein–coupled receptors (ADGRs). Representative conformations of the ADGRG2 receptor in the active, intermediate, and inactive states observed during dissociation of the IP15 peptide agonist dissociation are shown here.

