## Supplementary material for "Mechanisms of peptide agonist dissociation and deactivation of adhesion G-protein-coupled receptors": Table S1, Table S2, Supplementary Figures S1 to S5

**Protein-Protein Interaction-Gaussian accelerated Molecular Dynamics (PPI-GaMD)**

Based on GaMD, a new PPI-GaMD method [1] is developed for improved sampling of PPIs. Consider a system of ligand protein *L* binding to a target protein *P* in a biological environment *E*. The system comprises of *N* atoms with their coordinates $r\equiv\left\{ {\overset{⃑}{r}}_{1},\cdots,{\overset{⃑}{r}}_{N} \right\}$, and momenta $p\equiv\left\{ {\overset{⃑}{p}}_{1},\cdots,{\overset{⃑}{p}}_{N} \right\}$. The Hamiltonian of the system is formulated as:

$H\left( r, p \right)=K\left( p \right)+V\left( r \right)$,	(1)

where $K\left( p \right)$ and $V\left( r \right)$ represent the system’s kinetic and total potential energies, respectively. The potential energy term is divided into the following terms:

$V\left( r \right)=V_{P,b}\left( r_{P} \right)+V_{L,b}\left( r_{L} \right)+V_{E,b}\left( r_{E} \right)$

$$+ V_{PP,nb}\left( r_{P} \right)+V_{LL,nb}\left( r_{L} \right)+V_{EE,nb}\left( r_{E} \right)$$

$+V_{PL,nb}\left( r_{PL} \right)+V_{PE,nb}\left( r_{PE} \right)+V_{LE,nb}\left( r_{LE} \right)$	(2)

where $V_{P,b}$, $V_{L,b}$ and $V_{E,b}$ are the bonded potential energies in the protein *P*, protein *L* and environment *E*, respectively. Protein *P*, protein *L*, and environment *E*, each have self-non-bonded potential energies as $V_{PP,nb}$, $V_{LL,nb}$ and $V_{EE,nb}$ respectively*. P-L*, *P-E*, and *L-E's* related non-bonded interaction energies are $V_{PL,nb}$, $V_{PE,nb}$ and $V_{LE,nb}$ respectively. According to classical molecular mechanics force fields [2-3], the non-bonded potential energies are calculated as:

$V_{nb}=V_{elec}+V_{vdW}$,	(3)

where $V_{elec}$ and $V_{vdW}$ are the system electrostatic and van der Waals potential energies. The interaction energy between the protein binding partners is V_PL,nb_(r_PL_). In PPI-GaMD, we add boost potential selectively to the protein-protein interaction energy according to the GaMD method [4-6]:

${\Delta V}_{PL,nb}\left( r \right)=\left\{ \begin{aligned} \frac{1}{2}k_{PL,nb}\left( E_{PL,nb}-V_{PL,nb}\left( r_{PL} \right) \right)^{2}, &V_{PL,nb}\left( r_{PL} \right)<E_{PL,nb} \\ 0, &V_{PL,nb}\left( r_{PL} \right)\geq E_{PL,nb} \end{aligned} \right.$	(4)

where E*_PL,nb_* is the threshold energy for applying boost potential and *k_PL,nb_* is the harmonic constant. The PPI-GaMD simulation parameters are derived in the same manner as in the previous GaMD method [4-6]. When *E* is set to the lower bound as the system maximum potential energy (*E=V_max_*), the effective harmonic force constant$k_{0}$ can be formulated as:

$k_{0}=\min\left( 1.0, k_{0}^{'} \right)=min(1.0, \frac{\sigma_{0}}{\sigma_{V}}\frac{V_{max}-V_{min}}{V_{max}-V_{avg}})$,	(5)

where $V_{max}$, $V_{min}$, $V_{avg}$ and $\sigma_{V}$ are the maximum, minimum, average, and standard deviation of the boosted system potential energy, and $\sigma_{0}$ is the user-specified upper limit of the standard deviation of $\Delta V$ for proper reweighting [7]. The harmonic constant is calculated as $k=k_{0}\cdot\frac{1}{V_{max}-V_{min}}$ with ${0<k}_{0}\leq1$ . Alternatively, when the threshold energy *E* is set to its upper bound $E=V_{min}+\frac{1}{k}$,$k_{0}$ is set to:

$k_{0}=k_{0}^{"}\equiv(1-\frac{\sigma_{0}}{\sigma_{V}})\frac{V_{max}-V_{min}}{V_{avg}-V_{min}}$	(6)

if $k_{0}^{"}$ is found to be between *0* and *1*. Otherwise,$k_{0}$ is calculated using Eqn. (5).

In addition to selectively boosting the interaction energy between the protein partners *P* and *L*, another boost potential is applied on the remaining potential energy of the entire system to increase the conformational sampling and facilitate protein dissociation and rebinding. The second boost potential is calculated using the total system potential energy other than the interaction potential between the proteins as:

$\Delta V_{D}\left( r \right)=\left\{ \begin{aligned} \frac{1}{2}k_{D}\left( E_{D}-V_{D}\left( r \right) \right)^{2}, &V_{D}\left( r \right)<E_{D} \\ 0, &V_{D}\left( r \right)\geq E_{D} \end{aligned} \right.$	(7)

where *V_D_* is the total system potential energy other than the interaction potential between the proteins, E_D_ is the corresponding threshold energy for applying the second boost potential and *k_D_* is the harmonic constant, respectively. This provides us with the dual-boost PPI-GaMD with the total boost potential $\Delta V\left( r \right)=\Delta V_{PL,nb}\left( r_{PL} \right)+\Delta V_{D}\left( r \right)$.

**Energetic reweighting of PPI-GaMD for free energy calculations**

For energetic reweighting of PPI-GaMD simulations, the probability distribution along a selected reaction coordinate can be calculated as $p^{*}\left( A \right)$. Given the boost potential $\Delta V\left( r \right)$ of each frame in PPI-GaMD simulations, $p^{*}\left( A \right)$ can be reweighted to recover the canonical ensemble distribution, $p\left( A \right)$, as:

$p\left( A_{j} \right)=p^{*}\left( A_{j} \right)\frac{\left\langle e^{\beta\Delta V\left( \bar{r} \right)} \right\rangle_{j}}{\sum_{i=1}^{M} \left\langle{p^{*}\left( A_{i} \right)e}^{\beta\Delta V\left( \bar{r} \right)} \right\rangle_{i}}, j=1,\ldots, M$

(8)

where *M* is the number of bins, $\beta=k_{B}T$ and $\left\langle e^{\beta\Delta V\left( \bar{r} \right)} \right\rangle_{j}$ is the ensemble-averaged Boltzmann factor of $\Delta V\left( \bar{r} \right)$ for simulation frames found in the *j*^th^ bin. The ensemble-averaged reweighting factor can be approximated using cumulant expansion to reduce the energetic noise [4, 7]:

$\left\langle e^{\beta\Delta V\left( \bar{r} \right)} \right\rangle=exp\left\{ \sum_{k=1}^{\infty} \frac{\beta^{k}}{k!}C_{k} \right\}$

(9)

where the first two cumulants are given by:

$$C_{1}= \left\langle\Delta V \right\rangle,$$

$$C_{2}= \left\langle\Delta V^{2} \right\rangle-\left\langle\Delta V \right\rangle^{2}=\sigma_{\Delta V}^{2}.$$

(10)

The boost potential obtained from PPI-GaMD simulations usually shows near-Gaussian distribution [5]. Thus, cumulant expansion to the second order provides a more accurate reweighting [7]. The reweighted free energy $F\left( A \right)={-k}_{B}T\ln p\left( A \right)$ is calculated as:

$F\left( A \right)=F^{*}\left( A \right)-\sum_{k=1}^{2} \frac{\beta^{k}}{k!}C_{k}+F_{c}$

(11)

where $F^{*}\left( A \right)={-k}_{B}T\ln p^{*}\left( A \right)$ is the modified free energy obtained from PPI-GaMD simulation and $F_{c}$ is a constant.

**Kinetic reweighting of PPI-GaMD simulations**

In PPI-GaMD simulations, reweighting of protein binding kinetics followed a similar protocol using Kramer’s rate theory [6]. Given sufficient sampling of repetitive protein dissociation and binding in the simulations, we recorded the time periods for the protein sampled in the bound (τ*_B_*) and unbound (τ*_U_*) states. The protein binding and dissociation rate constants (*k*_off_ and *k*_on_) were calculated as:

$k_{off}=\frac{1}{\tau_{B}}$.	(12)

$k_{on}=\frac{1}{\tau_{U} [L]}$,	(13)

where [L] is the ligand protein concentration

Based on Kramers' rate theory, the rate of a chemical reaction in the large viscosity limit is determined by [6]:

$k_{R}\cong\frac{w_{m}w_{b}}{2\pi\xi}e^{-{\Delta F}/{k_{B}T}}$,	(14)

where $w_{m}$ and $w_{b}$ are frequencies of the approximated harmonic oscillators (also referred to as curvatures of free energy surface) near the energy minimum and barrier, respectively, $\xi$ is the frictional rate constant and $\Delta F$ is the free energy barrier of transition. The friction constant $\xi$ is related to the diffusion coefficient *D* with $\xi=k_{B}T/D$. The apparent diffusion coefficient *D* is obtained by dividing the kinetic rate calculated using the transition time series obtained directly from simulations by the probability density solution of the Smoluchowski equation [8]. In order to reweight protein kinetics from the simulations using Kramer’s rate theory, the free energy barriers of protein binding and dissociation are calculated from the original (reweighted, ***∆F***) and modified (no reweighting, ***∆F****) PMF profiles, similarly for curvatures of the reweighed (*w*) and modified ($w^{*}$, no reweighting) PMF profiles near the protein bound (“B”) and unbound (“U”) low-energy wells and the energy barrier (“Br”), and the ratio of apparent diffusion coefficients from simulations without reweighting (modified, $D^{*}$) and with reweighting(*D*). The resulting numbers are then plugged into Eq. (14) to obtain accelerations of the protein binding and dissociation rates during the PPI-GaMD simulations, which allows us to obtain the original kinetic rate constants [6].

**Table S1.** **Summary of the PPI-GaMD simulations performed on the human ADGRG2 in the presence of the IP15 peptide agonist.**

| **System** | **System Size** | **ID** | **Simulation length** | **Boost Potential (kcal/mol)** |
| --- | --- | --- | --- | --- |
| **IP15-ADGRG2** | 79,721 atoms | Sim1 | 1500ns | 30.08 ± 4.40 |
|  |  | Sim2 |  | 28.85 ± 4.55 |
|  |  | Sim3 |  | 30.99 ± 5.41 |
|  |  | Sim4 |  | 29.78 ± 4.88 |
|  |  | Sim5 |  | 28.57 ± 4.62 |
|  |  | Sim6 |  | 31.03 ± 5.38 |
|  |  | Sim7 |  | 31.68 ± 4.73 |
|  |  | Sim8 |  | 30.61 ± 5.42 |

**Table S2.** **Summary of the PPI-GaMD simulations performed on the human ADGRG1 in the presence of the P7 peptide agonist.**

| **System** | **System Size** | **ID** | **Simulation length** | **Boost Potential (kcal/mol)** |
| --- | --- | --- | --- | --- |
| **P7-ADGRG1** | 74,044 atoms | Sim1 | 1500ns | 49.17 ± 5.62 |
|  |  | Sim2 |  | 46.34 ± 5.01 |
|  |  | Sim3 |  | 48.53 ± 4.80 |
|  |  | Sim4 |  | 45.52 ± 6.64 |
|  |  | Sim5 |  | 46.78 ± 4.12 |
|  |  | Sim6 |  | 46.82 ± 4.22 |
|  |  | Sim7 |  | 47.12 ± 5.31 |
|  |  | Sim8 |  | 48.22 ± 4.80 |

**Figure S1. (A)** Simulation starting structure of the ADGRG2-IP15 complex with the G protein removed **(B)** Computational model of the ADGRG2-IP15 system embedded in membrane lipids and solvated in aqueous medium. The phosphatidylcholine (POPC) membrane lipids were rendered as cyan sticks, the ADGRG2 receptor as grey cartoons, and the sodium and chlorine ions as yellow and green spheres.

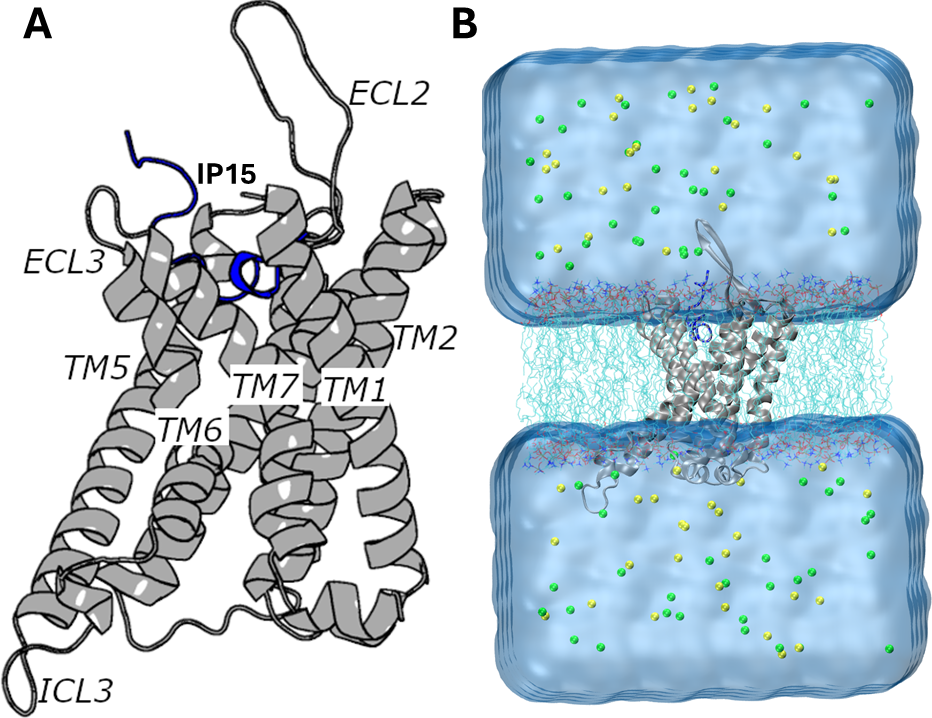

**Figure S2. (A)** The low-energy conformation of the IP15-ADGRG2 complex in the “Bound/Active” (B/A) state compared with the cryo-EM structure (grey, PDB: 7WUI). **(B)** Critical interactions between IP15 (blue) and ADGRG2 (grey) observed in the B/A state. The peptide agonist formed hydrogen-bonding (yellow dash lines) and hydrophobic interactions with receptor residues Y^1.44^, F^2.64^, W779^ECL2^, Y^5.36^, F^6.57^ and F^7.42^.

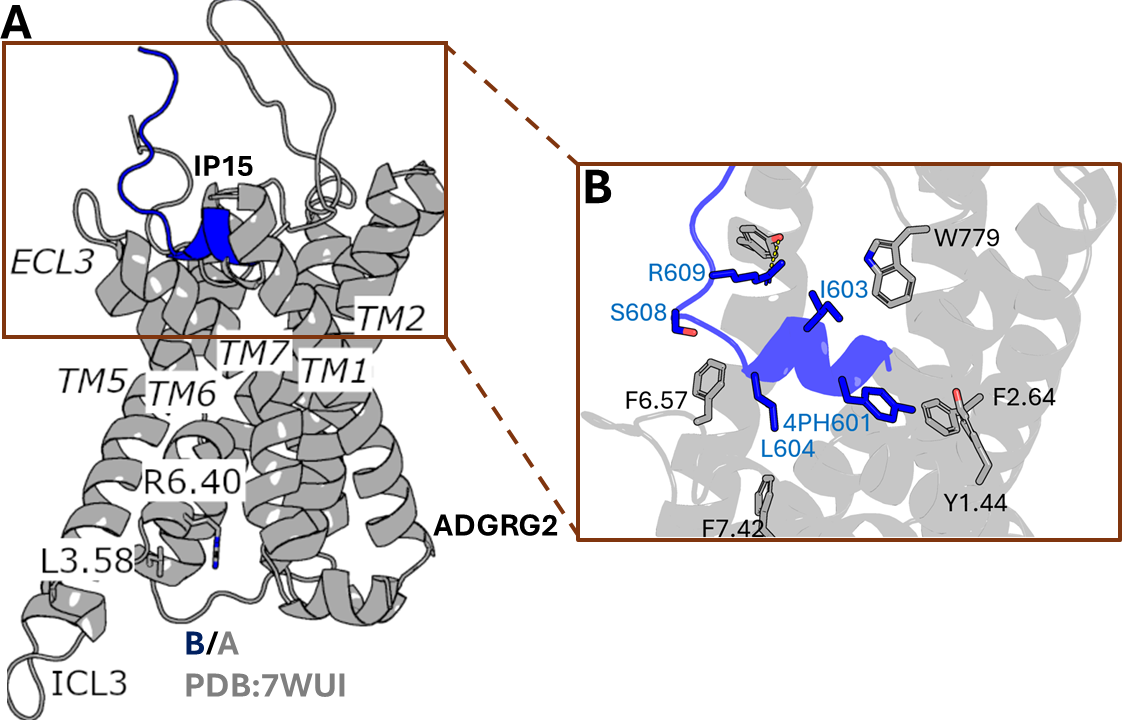

**Figure S3. (A)** 2D potential of mean force (PMF) free energy profile of the distance between residues W779^ECL2^-W^6.60^(ECL3) and the IP15 RMSD calculated by combining the eight PPI-GaMD simulations. The low-energy states are labeled as “Bound/Closed” (B/Closed), “Partially Dissociated 2/Open” (PD2/Open) and “Unbound/Open” (U/Open). **(B)** Opening of the ECL3 in the “PD2/Open” state compared with the “B/Closed” state.

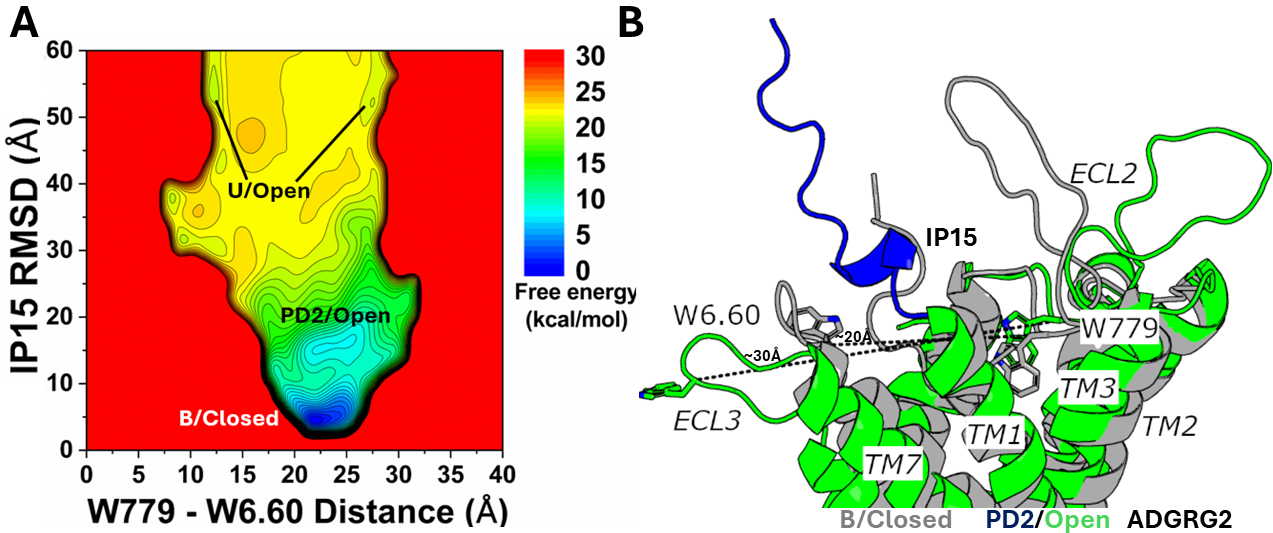

**Figure S4. (A)** Simulation starting structure of the ADGRG1-P7 complex with the G protein removed. **(B)** Computational model of the ADGRG1-P7 system embedded in membrane lipids and solvated in aqueous medium. The phosphatidylcholine (POPC) membrane lipids were rendered as cyan sticks, the ADGRG1 receptor as grey cartoons, and the sodium and chlorine ions as yellow and green spheres.

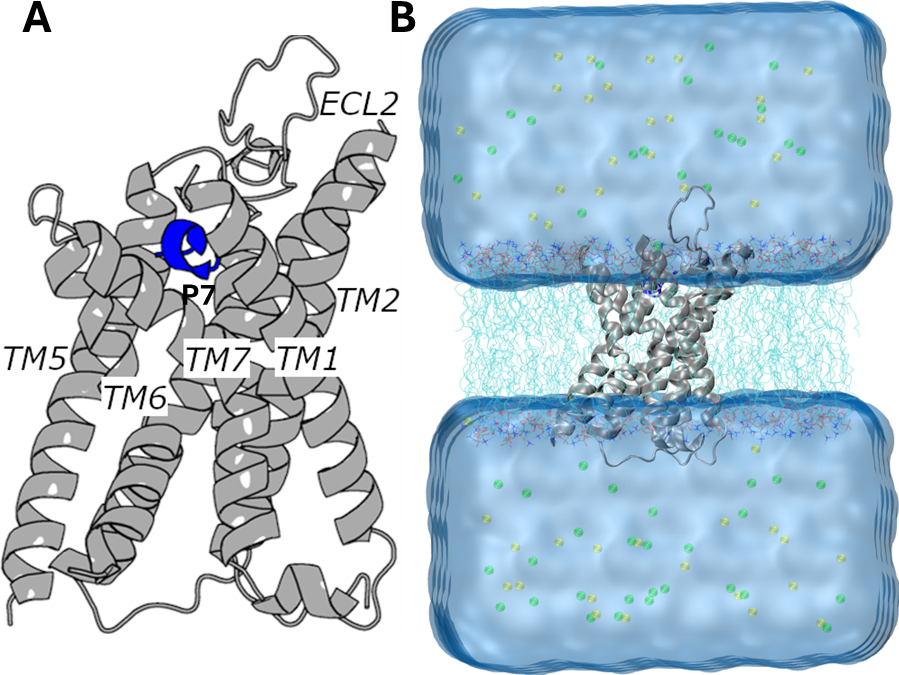

**Figure S5. (A)** The low-energy conformation of the P7-ADGRG1 complex in the “Bound/Active” (B/A) state compared with the cryo-EM structure (grey, PDB: 7SF8). **(B)** Critical interactions between P7 (blue) and ADGRG1 (grey) observed in the B/A state. The peptide agonist formed hydrophobic interactions with receptor residues F^2.58^, W563^ECL2^, Y^5.36^, F630^ECL3^, W^6.53^ and F^7.42^.

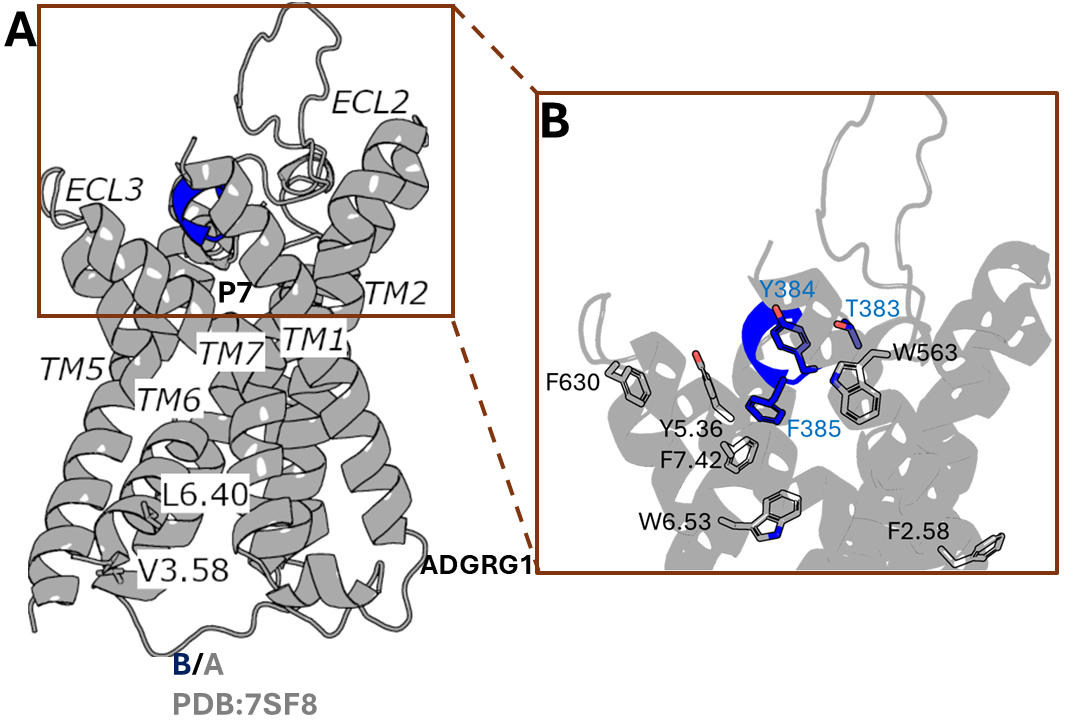
