## Supplementary figures and images for "Mechanisms of peptide agonist dissociation and deactivation of adhesion G-protein-coupled receptors"

### Supplementary Figure S1

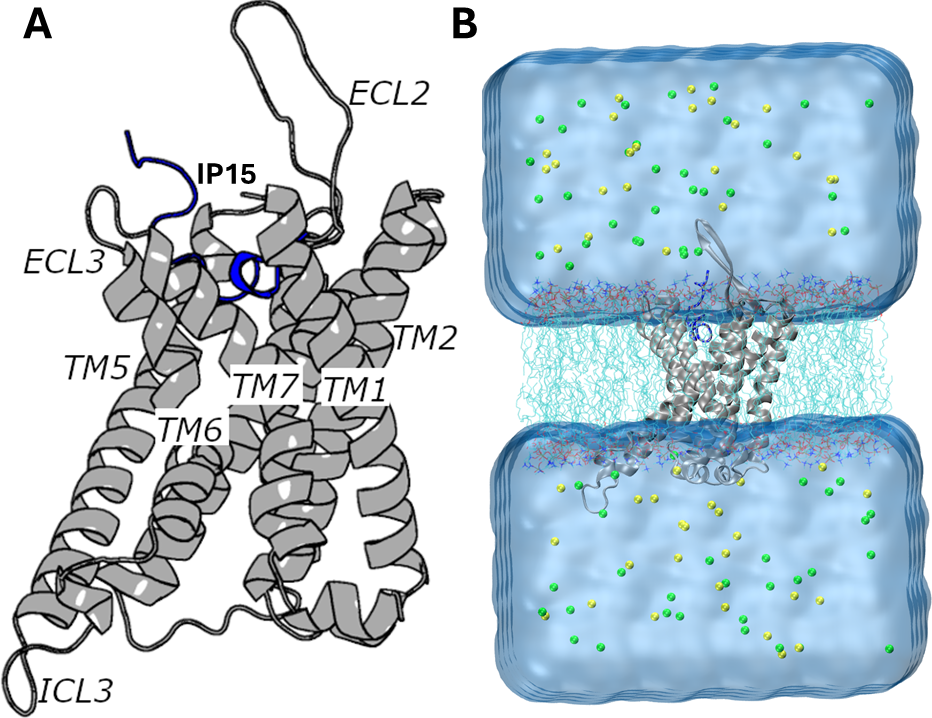

### Supplementary Figure S2

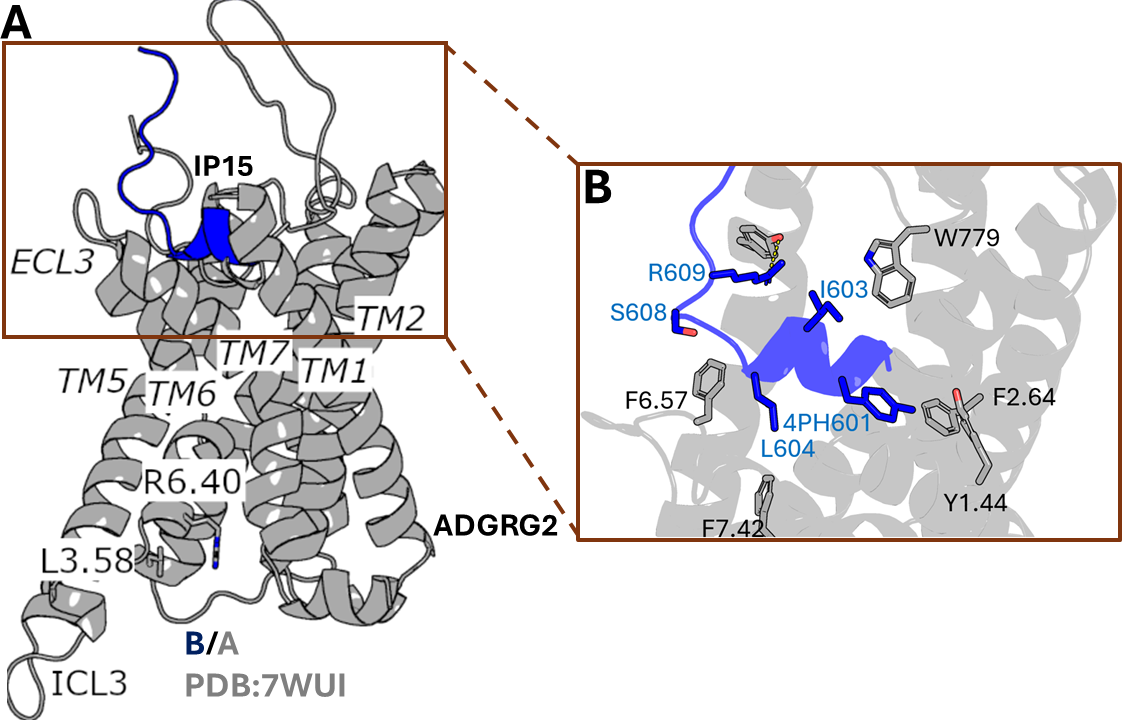

### Supplementary Figure S3

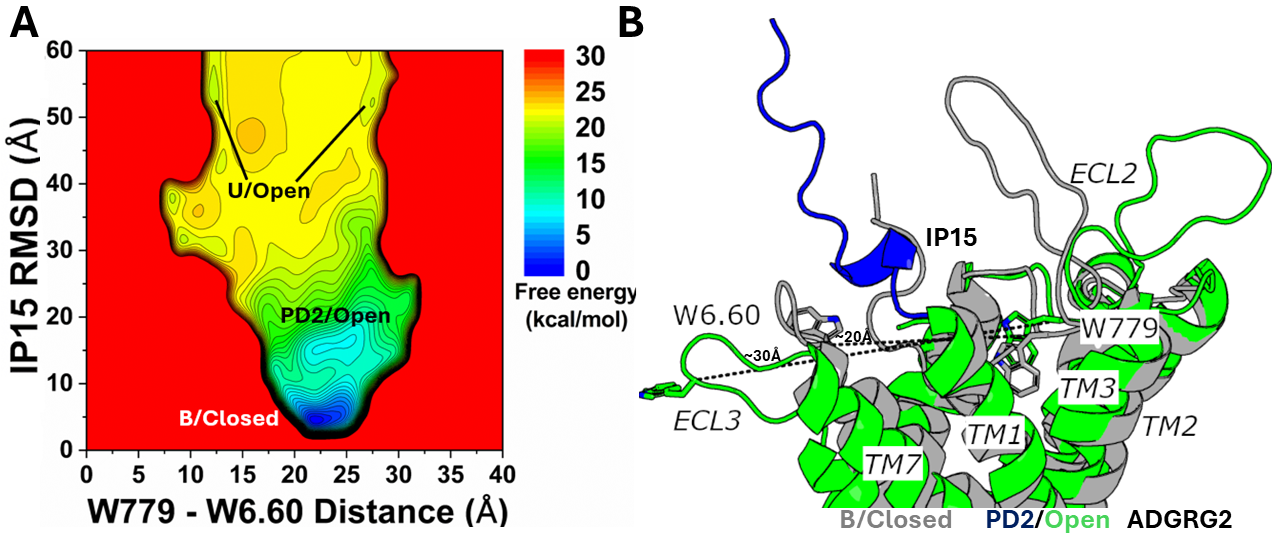

### Supplementary Figure S4

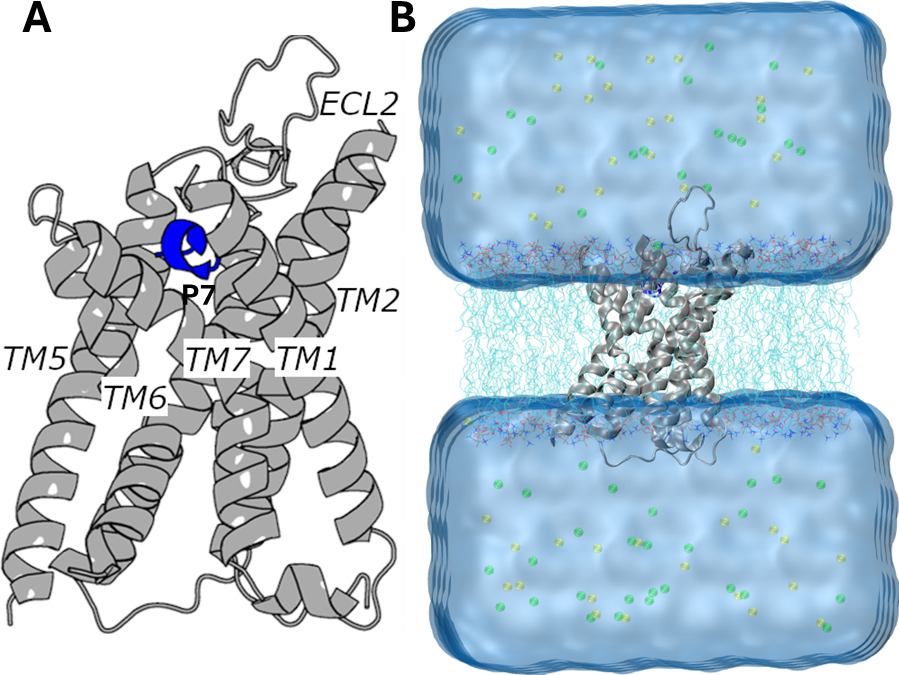

### Supplementary Figure S5

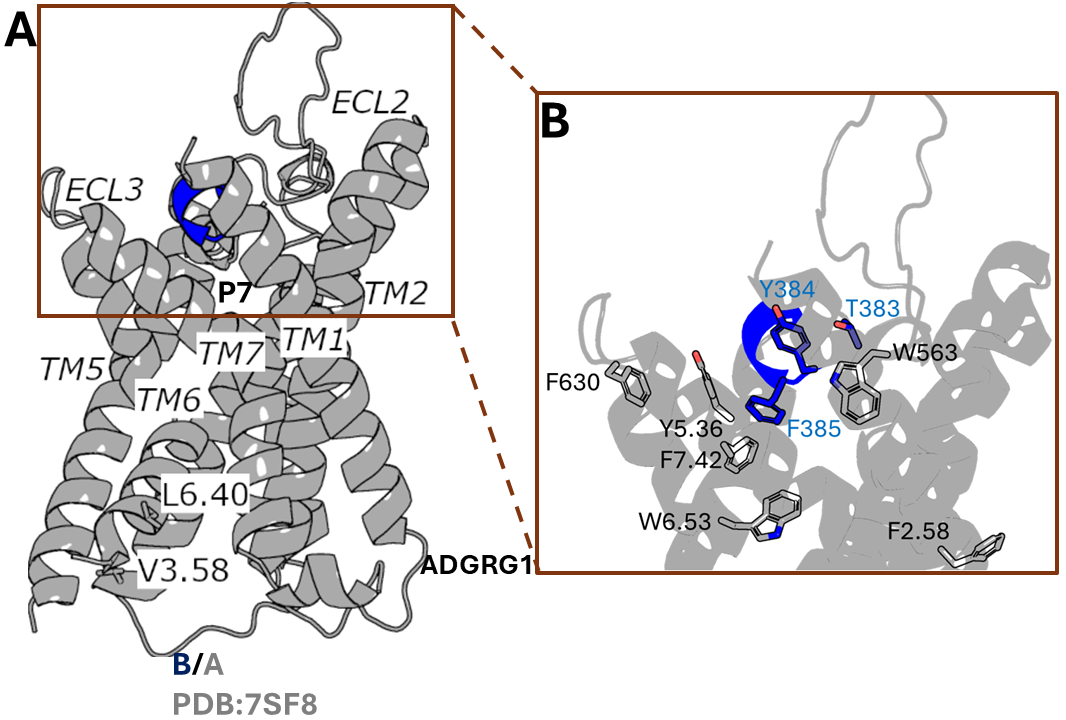
